# Ectopical expression of bacterial collagen-like protein supports its role as adhesin in host-parasite coevolution

**DOI:** 10.1101/2023.07.14.549037

**Authors:** Benjamin Huessy, Dirk Bumann, Dieter Ebert

## Abstract

For a profound understanding of the mechanisms of antagonistic coevolution, it is necessary to identify the coevolving genes. The spore-forming bacterium *Pasteuria ramosa* and its host, the microcrustacean *Daphnia*, are a well-characterized paradigm for co-evolution, but the underlying genes remain largely unknown. A genome-wide association study identified a polymorphic carboxy-terminal globular domain of *Pasteuria* collagen-like protein 7 (Pcl7) as a candidate mediating parasite attachment and driving its coevolution with the host. Since *P. ramosa* cannot currently be genetically manipulated, we used *Bacillus thuringiensis* as a surrogate parasite to express a fusion protein of a Pcl7 carboxy-terminus from *P. ramosa* and the amino-terminal domain of a *B. thuringiensis* collagen-like protein. Mutant *B. thuringiensis* (Pcl7-*Bt*) spores but not wild-type *B. thuringiensis* (WT-*Bt*) spores, attached to the same site of susceptible hosts as *P. ramosa*. Furthermore, Pcl7-*Bt* spores attached readily to host genotypes that were susceptible to the *P. ramosa* clone that was the origin of the Pcl7 C-terminus, but only slightly to resistant host genotypes. These findings indicated that the fusion protein was properly expressed and folded and demonstrated that indeed the C-terminus of Pcl7 mediates attachment in a host genotype-specific manner. These results provide strong evidence for the involvement of a CLP in the coevolution of *Daphnia* and *P. ramosa* and opens new avenues for genetic epidemiological studies of host-parasite interactions.

**150-word “Importance” paragraph:** During host-parasite coevolution, hosts evolve to evade the damaging effect of the parasite, while parasites evolve to maximize their benefits by exploiting the host. The genes underlying this coevolution remain largely unknown. For the prime model-system for coevolutionary research, the crustacean *Daphnia* and the parasite *Pasteuria ramosa*, collagen-like proteins (CLPs) in *Pasteuria* were suggested to play a crucial role for host-parasite interactions. Here we report that transferring part of a CLP coding gene from the unculturable *P. ramosa* to *Bacillus thuringiensis* (*Bt*), confirmed the function of this protein as a genotype-specific adhesin to the host’s cuticle. Our finding highlights the importance of a CLP in host-parasite interactions and will enable us to explore the population genetic dynamics of coevolution in this system.

## Introduction

Antagonistic coevolution between hosts and parasites has been suggested to be a major driver in evolution, presumably underlying diverse biological phenomena, such as the extraordinary genetic variation at the major histocompatibility complex (MHC) of jawed vertebrates and R-genes in plants, the parasite hypothesis about the evolution of sexual selection, the evolution of genetic recombination and the evolution of immune systems (1–3). While great progress has been made in our understanding of coevolution and its consequences at the phenotypic level, much less is known about the underlying genetics (4–6). However, current theories about the mechanism of coevolution are genetic models, such as the selective sweep model, where new beneficial mutations sweep to fixation in both antagonists, and balancing selection models, where alleles at specific loci interact in a manner that generates negative frequency dependent selection (7, 8). Therefore, to test these models and understand the evolutionary dynamics during coevolution, we need to identify the genes involved in host-parasite interactions. This is particularity challenging in non-model systems, where genetic tools are largely lacking.

A prime model system for coevolution research is the water flea *Daphnia* and the bacterial parasite *Pasteuria ramosa. P. ramosa* is a highly virulent obligate parasite of its planktonic crustacean host. It cannot be cultured outside its host. For the *Pasteuria–Daphnia* system, coevolutionary dynamics have been demonstrated in natural and experimental settings, with negative frequency dependent selection being the main explanation for the observed dynamics (9–11). The system is renowned for its strong genotypic infection specificity (12–14), but the genes responsible for this specificity have not been identified, even so candidates have been suggested for both host and parasite (15–17). During infection, dormant *P. ramosa* endospores are taken up by the filter-feeding *Daphnia* and shed their exosporium, revealing numerous peripheral fibres (13). These activated spores attach to the cuticles of susceptible *Daphnia*, most commonly to the oesophagus or the hindgut wall (15, 18, 19). The current understanding is that the peripheral fibres of the activated spores may be collagen-like proteins (CLPs) that act as adhesins on surface components on the *Daphnia* epithelium.

Collagen-like proteins (CLPs), proteins with high similarity to eukaryotic collagens, have been identified in a range of prokaryotes (20–22) including human-pathogenic species such as *Bacillus anthracis* (23), *Legionella pneumophila* (24) and several *Streptococcus* species (25). CLPs typically attach to cell walls (26, 27) and contain a rod-shaped collagenous domain near the cell surface (28–30). Research on bacterial CLPs indicates that they play a pivotal role in host–pathogen interactions during the initial stages of infection for attachment to host cells and surfaces (31–36). While most bacteria contain only a few genes that encode for CLPs (21), the endospore-forming bacteria of the Gram-positive *Pasteuria* genus carries up to 50 CLP-encoding genes (20, 37), one of them, Pcl7 of *P. ramosa*, was suggested to be responsible for the highly specific interaction with the host and may play an important role in their coevolution (19). Conducting a genome-wide-association study Andras et al. (19) discovered multiple phenotype-associated sequence polymorphisms in the *P. ramosa pcl7* gene encoding for a *Pasteuria* collagen-like protein,. In its C-terminal domain (CTD), *pcl7* contains seven single-nucleotide sequence polymorphisms that correlate perfectly with infection phenotype and that encompass considerable changes in the size, hydrophobicity, and charge of the respective amino acids. These findings suggest that Pcl7 may be crucial for spore–host attachment and, furthermore, that sequence variation in Pcl7 may be important for determining the high specificity of the bacterial spore’s adhesion to the host epithelium in the oesophagus. The aim of this study was to test the hypothesis that *pcl7* is responsible for attachment to the host’s oesophagus through experimental manipulation of Pcl7.

Genetic engineering on *P. ramosa* has not yet been successful because its rigid exosporium resists lysis and degradation (38), preventing us from generating *pcl7* mutants. However, CLPs can be engineered in the *Bacillus cereus* group (39–43), providing a heterologous system for studying *pcl7* and other *Pasteuria* factors (44). BclA, a structural homolog of Pcl7 (19) is part of the exosporium in the entire *B. cereus* group including *B. thuringiensis*, a species with well-developed techniques for laboratory experiments.

Members of the *B. cereus* group develop spores encapsulated by an exosporium composed of two defined layers (29): a primary basal layer and an outermost hair-like structure (45, 46). BclA makes up the majority of this hair-like structure (23). It is expressed at the spore surface late during sporulation and requires the specific amino acid sequence motif “LVGPTLPPIPP” for incorporation into the exosporium (47).

Here, we used the collagen-part of BclA as a display platform for the CTD of Pcl7 in *B. thuringiensis* to obtain a surrogate parasite that displays the key part of Pcl7 on its surface. Analogous display systems have already been used to express functional fusion proteins (48–50). We showed that the Pcl7 CTD mediated attachment of the surrogate parasite spores to the oesophagus wall of *D. magna* and that this single protein part was sufficient to recapitulate host genotype specificity of the donor *P. ramosa* clone. These data demonstrate the key role of Pcl7 CTD in this paradigmatic host-pathogen system.

## Material and Methods

**Table 1.** Material used in the study.

A. *Daphnia magna* genotypes (=clones) used in the study
| Clone | Country of Origin (collection site) | C1 Resistotype |
| --- | --- | --- |
| HU-HO-2 | Hungary | Susceptible |
| NO-AA-1 | Norway | Susceptible |
| RU-AST1-1 | Russia | Susceptible |
| CH-H-2016-h-34 | Switzerland | Susceptible |
| FI-Xinb3 | Finland | Resistant |
| DE-G1-106 | Germany | Resistant |

B. Bacterial strains and plasmids used in this study
| Strain or plasmid | Characteristics | Reference |
| --- | --- | --- |
| **Bacterial strain** |  |  |
| *B. thuringiensis* 407 Cry | Acrystalliferous *B. thuringiensis* strain | (52) |
| *E. coli* DC10B | *mcrA* $\Delta(mrr-hsdRMS-mcrBC)$ $\phi80lacZ\Delta M15 \Delta lacX74 recA1 araD139 \Delta(ara-leu)7697 galU galK rpsL endA1 nupG \Delta dcm$ | (53) |
| *P. ramosa* C1 and C19 | cloned lines of *P. ramosa*, isolated from natural populations | (13) |
| **Plasmid** |  |  |
| pBKJ223 | Plasmid encoded homing restriction enzyme I-SceI, promoting the second homologous recombination event in procedure for making markerless deletion mutant, Tet<sup>r</sup> | (54) |
| pMAD-I-SceI | Plasmid used for making deletion mutants, containing I-SceI restriction site that can be cleaved by the homing endonuclease I-SceI, Apr, Eryr | (55) |

### Media

*E. coli* DC10B was cultured in lysogenic broth (LB) (10 g/L bacto tryptone, 5 g/L yeast extract, 10 g/L NaCl) medium. *B. thuringiensis* was cultured in tryptic soy broth (TSB) (30 g/L bacto tryptic soy broth) medium. LB-lowsalt (LB-ls) (10 g/L bacto tryptone, 5 g/L yeast extract, 5 g/L NaCl) was used to prepare electrocompetent cells. For electroporation, Super-Optimal broth with Catabolite Repression (SOC) (20 g/L tryptone, 5 g/L yeast extract, 0.5 g/L NaCl, 4.8 g/L MgSO4, 0.186 g/L KCl, and 3.6 g/L glucose) was used. For selection, we used LB agar plates (10 g/L bacto tryptone, 5 g/L yeast extract, 10 g/L NaCl, 15 g/L agar) and TSB agar plates (30 g/L bacto tryptic soy broth, 15 g/L agar) with erythromycin at a final concentration of 5 μg/mL, ampicillin at a final concentration of 100 μg/mL, and anhydrous tetracycline at a final concentration of 8 μg/mL.

### Preparation of electrocompetent cells and electroporation

To prepare electrocompetent cells, 100 mL of fresh LB-ls was inoculated at OD_600nm_ 0.01 from an overnight culture and grown to OD_600nm_ 0.08 at 37 °C, 180 rpm. The culture was distributed to two 50-mL falcon tubes and put on ice for 20 min. The tubes were spun down at 4 °C, 8000 rpm for 8 min, and the supernatant was removed. The pellets were then washed with 30 mL cold sterile water containing 15 % glycerol (AppliChem, A1123,1000), and the tubes were spun down at 4 °C, 8000 rpm for 8 min. This washing step was repeated twice. The supernatant was discarded, and the pellet was suspended in 1 mL cold sterile water containing 15 % glycerol. Finally, 100 μL aliquots were stored at -80 °C or used directly for electroporation.

For electroporation of plasmid DNA, DNA (100 ng) was added to thawed, electrocompetent cells (100 μL) in a 1-mm electroporation cuvette on ice. For *E. coli* DC10B, a single pulse at 1.8 V (Gene Pulser Xcell Electroporation Systems, Bio-Rad Laboratories) was applied. For *B. thuringiensis*, a single pulse at 2.5 V was applied. Immediately after the pulse, 900 μL of pre-warmed SOC media was added, and the entire volume was transferred to a fresh 1.5-mL Eppendorf tube. The tube was incubated and shaken at 37 °C, 180 rpm for 1 h. Samples were spun down for 4 min at 11000 rpm. We removed 900 μL of supernatant and plated the remaining volume (100 μL) on LB agar containing the respective antibiotics. Plates were incubated overnight at 37 °C.

### Isolation of genomic DNA

Genomic DNA was isolated using a DNeasy Blood & Tissue Kit (Qiagen GmbH, Hilden, Germany). A 2-mL overnight culture of *B. thuringiensis* in TSB media was spun down at 8.000 rpm for 4 min. The supernatant was discarded, and the pellet was resuspended in 200 μL AL Buffer and 20 μL of proteinase K. The sample was incubated at 56 °C, 500 rpm for 10 min, and 200 μL >99 % ethanol was added to it. It was then mixed thoroughly, transferred to a DNeasy Mini spin column, and spun down at 10000 rpm for 1 min; the flow through was discarded. 500 μL of Buffer AW1 was then added and spun down at 10000 rpm for 1 min, and the flow through was discarded. Finally, 500 μL of Buffer AW2 was added and spun down at 13200 rpm for 1 min. The column was transferred to a fresh 1.5-mL Eppendorf tube, and the DNA was eluted with 50 μL sterile water, incubated at room temperature for 1 min and centrifuged at 10000 rpm for 30 sec. The concentration of DNA was measured using a Colibri Microvolume Spectrometer (Berthold Technologies, Bad Wildbad, Germany).

### Molecular Biology

Plasmids were constructed by Gibson Assembly (56). Flanking regions (∼750 bp) of the target locus were PCR-amplified from genomic DNA (with primers 1 & 2 as well as 5 & 6, Table 2) using Phanta Max Super-Fidelity DNA Polymerase (Vazyme Biotech, Nanjing, China), and primers were designed with SnapGene (v. 5.2, Gibson assembly tool). The 501 bp C-terminal *pcl7* sequence was synthesized (LubioScience GmbH, Zurich, Switzerland), and PCR-amplified using primers 3 & 4. The pMAD-I-SceI vector was amplified with primers 7 & 8 in a long-range PCR. The resulting three fragments and the vector were fused using the Hifi DNA Assembly Master Mix (New England Biolabs, Ipswich, USA) at 50 °C for 1 h. The reaction mix contained approximately 40 ng of each fragment and 150 ng of vector for a total volume of 20 μL. *E. coli* DC10B was transformed with the resulting product. Plasmid DNA was purified from an overnight culture using a plasmid miniprep kit (ZymoPURE, ZymoResearch). Sequence-verified plasmids were electroporated into *B. thuringiensis*. Cells were incubated at 28 °C for 1 h and plated on TSB agar containing 5 μg/mL erythromycin.

**Table 2.**
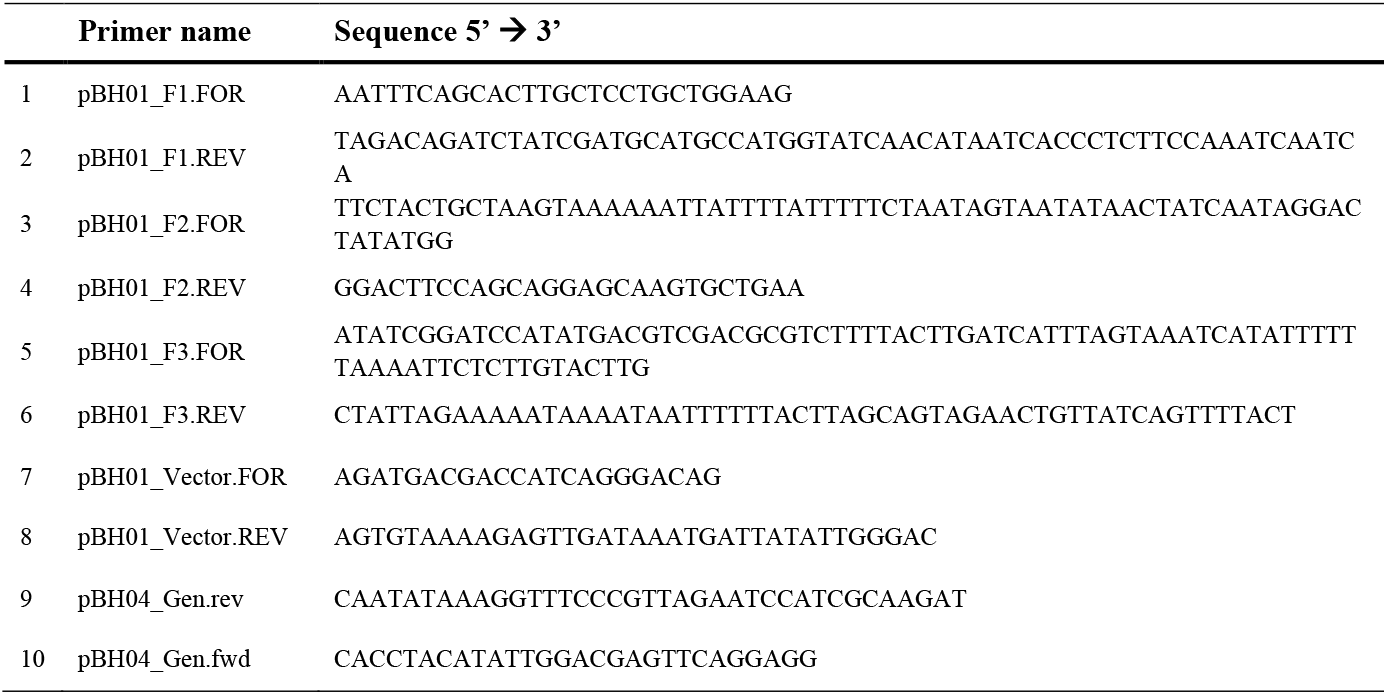
Primers used in this study.

The allelic-exchange procedure was done as described (54) with some modifications. *B. thuringiensis* integrants were isolated by shifting transformants obtained by electroporation from 28 °C to 37 °C in the presence of 5 μg/mL erythromycin. pMAD-I-SceI cannot replicate in *B. thuringiensis* at 37 °C, so only cells which integrated the plasmid into the chromosome could propagate. Colonies were screened for the orientation of their insert using PCR (with primers 9 & 10, Table 2). Colonies that harboured the right-hand insert were pooled and inoculated in 4 mL fresh TSB medium containing 5 μg/mL erythromycin at 37 °C, 180 rpm overnight. The overnight culture was diluted 1:100 in 50 mL fresh TSB medium containing 5 μg/mL erythromycin and incubated at 37 °C, 180 rpm to an OD_600nm_ of 0.8 to prepare electrocompetent cells. The cells were transformed with pBKJ223 and plated on TSB agar containing 8 μg/mL tetracycline. Colonies were pooled and inoculated in 4 mL fresh TSB medium containing 8 μg/mL tetracycline at 37 °C, 180 rpm for 6 h. Serial dilutions (10^−1^, 10^−2^, 10^−3^, 10^−4^) were prepared and 100 μL of each dilution was plated on TSB agar plates containing 8 μg/mL tetracycline. Single colonies were patched on TSB agar with 8 μg/mL tetracycline as well as TSB agar with 5 μg/mL erythromycin to screen for loss of erythromycin-resistance. Erythromycin-sensitive clones were picked and stored in 50 μL LB-Glycerol (15 %) at -20 °C. The clones were screened for the desired insert (Fig 1C; recombination through homologous regions “Y”) using primers 9 & 10. Clones with confirmed allelic exchange were inoculated into 4 mL fresh TSB media and incubated at 37 °C, 180 rpm overnight. Genomic DNA was prepared and Sanger-sequenced (Microsynth AG, Balgach, Switzerland). Correct clones were inoculated in 4 mL fresh TSB and incubated at 37 °C, 180 rpm overnight. The next day freezer stocks were generated and stored at -80 °C for later use.

**Figure 1.**
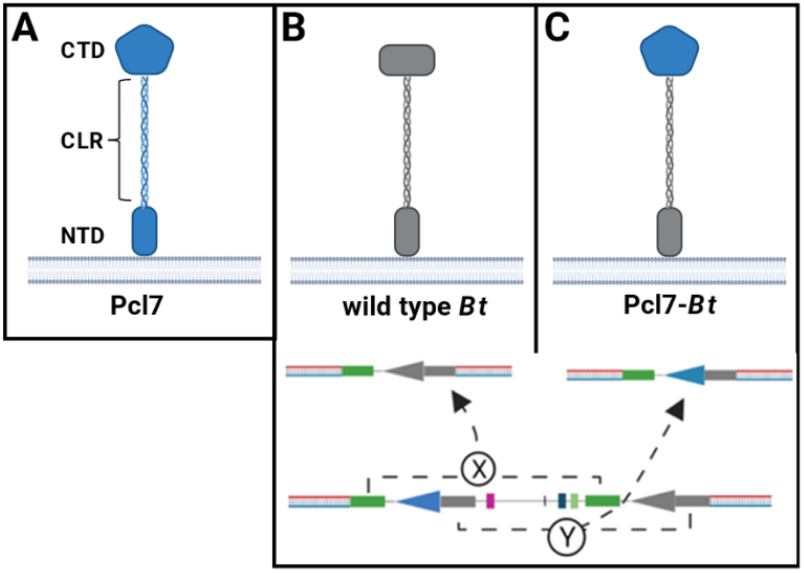
Graphic overview of the construction of the Pcl7-*Bt* fusion. **(A)** Pcl7 consists of three domains: an amino-terminal (NTD), a central collagen-like region (CLR) and a carboxy-terminal globular domain (CTD). **(B)** Homologous recombination can occur either through homologous regions “X”, resulting in the regeneration of the wild type gene and protein, **(C)** or through “Y”, resulting in the generation of the Pcl7-*Bt* fusion. Figure created with Biorender.com.

### Sporulation and purification of spores

Spores were purified as described by (57) with some modifications. An overnight culture of *B. thuringiensis* in 4 mL of TSB was grown at 37 °C, 180 rpm. The overnight culture was diluted 1:100 in 50 mL fresh TSB media containing 0.1 mmol MnSO_4_ (PanReac, AppliChem) and grown at 37 °C, 180 rpm for 10 days. Cultures were transferred to a 50-ml falcon tube and stored on ice for 20 min. The tubes were spun down at 4 °C, 10000 rpm for 10 min. The pellet was suspended in 20 mL of 50 mM Tris-HCL (ph 7.2; PanReac, AppliChem) with an addition of 50 μg/mL lysozyme (from hen egg whites, Fluka).

Samples were incubated at 37 °C, 180 rpm for 1 h. The sample was spun down at 4 °C, 10000 rpm for 10 min, and the pellet was suspended in cold sterile water. This washing step was repeated twice. The pellet was then suspended in 5 mL 0.05 % SDS solution (Sigma, D6750-10G) by vortexing and incubated for 5 min at room temperature. The sample was then spun down at 4 °C, 10000 rpm for 10 min, and the pellet was suspended in cold sterile water. This washing step was repeated five times. After the final washing step, the pellet was suspended in 5 mL of cold, sterile water and stored at 4 °C for later use. Spores were counted using a Neubauer improved chamber (Paul Marienfeld GmbH, Lauda-Konigshofen, Germany) with a chamber depth of 0.1 mm.

### Fluorescent labelling of spores

*Pasteuria ramosa* spores were isolated by homogenizing infected *Daphnia* in ADaM followed by centrifugation at room temperature, 8000 rpm for 5 min. *Bacillus thuringiensis* spores were thawed and centrifuged at room temperature, 8000 rpm for 5 min. *P. ramosa* or *B. thuringiensis* pellets were suspended in 0.8 mL of 0.1 M sodium bicarbonate (pH 9.1; Sigma, S5761-500G) with 2 mg/mL of fluorescein-5(6)-isothiocyanate (Sigma-Aldrich, Miss, USA). The samples were then incubated in the dark at room temperature, 1600 rpm shaking for 2 h, followed by centrifugation at room temperature, 8000 rpm for 4 min. The pellet was suspended in 0.8 mL sterile water and centrifuged at room temperature, 8000 rpm for 4 min; the supernatant was then removed. This washing step was repeated three times. Spore suspensions were stored in sterile water at 4 °C in the dark for further use.

### Attachment assay

*Daphnia* were individually placed into a 96 well plates containing 150 μL of ADaM per well. 10 μL of spore solution containing **∼** 500 fluorescently labelled *P. ramosa* spores were added to each well and incubated in the dark for 5 min. For *B. thuringiensis*, 10 μL of spore solution containing **∼** 50’000 labelled *B. thuringiensis* spores were added to each well and incubated in the dark for 5 min. The entire liquid volume in each well was removed and replaced with 150 μL fresh ADaM. This washing step was repeated twice, after which the entire liquid volume in each well was removed. The *Daphnia* were placed individually on a microscopy slide using a toothpick. A glass cover slide was applied to the *Daphnia* gently to avoid crushing it. Extended focus images were taken using Leica Application Suite (v. 4.12, using package “montage”) with a Leica DM6 B (Leica Microsystems, Wetzlar, Germany) microscope fitted with a Leica DFC 7000T camera and a GFP Filter cube (Excitation Filter BP 470/40).

### Competitive attachment assay

*Daphnia* were individually placed into a 96-well plate containing 150 μL of ADaM per well. We then added 10 μL of spore solution to each well according to each treatment—for C1/C19: 50 labelled *P. ramosa* spores; for WT-*Bt* + C1/C19: **∼** 20’000 labelled *B. thuringiensis* spores followed by 50 labelled *P. ramosa* spores; and for Pcl7-*Bt* + C1/C19: **∼** 20’000 labelled Pcl7-*Bt* spores followed by 50 labelled *P. ramosa* spores—and incubated them in the dark for 5 min. The entire liquid volume in each well was removed and replaced with 150 μL fresh ADaM. This washing step was repeated twice to remove excess spores. The *Daphnia* were then placed individually on a microscopy slide and a glass cover slide was gently applied to the *Daphnia* to avoid crushing them. The number of *P. ramosa* spores attached to the *D. magna* oesophagus were then counted by manually scanning the z-stacks.

### Selection of candidate collagen-like proteins

The N-terminal domain and collagen-like region of three out of eight *bclA* homologs in *B. thuringiensis* (Table 3) were selected based on their similar length and sequence as *pcl7* to construct fusions with the *pcl7* CTD.

**Table 3.** Candidate collagen-like proteins in *B. thuringiensis*. Three candidate *B. thuringiensis* genes with similar N-terminal domains and amino acid sequences as *pcl7*. Deviations from the specific sequence motif, likely required for incorporation into the exosporium, are marked in bold. Sequences were obtained from GenBank (58)(accession no. CP003889).

| Locus tag | Product | Reasoning | Motif |
| --- | --- | --- | --- |
| BTB_c12600 | Collagen-like exosporium surface protein | Short and highly similar NTD to *pcl7* | LVGPTLPPIPP |
| BTB_c35490 | Collagen triple helix repeat protein | Similar NTD to *pcl7* | LIGPTLPSIPP |
| BTB_c38740 | Triple helix repeat-containing collagen | Similar NTD to *pcl7* | IIGPTLPPVPP |

## Results

To validate the role of the C-terminal domain (CTD) of the collagen-like protein Pcl7 of *Pasteuria ramosa* as an adhesin that recognizes the host *Daphnia magna* in a genotype-specific manner, we employed surface presentation on *Bacillus thuringiensis* spores. Specifically, we fused the Pcl7-CTD to the N-terminal domain of the main collagen-like exosporium surface protein of *B. thuringiensis* (BTB_c12600, Table 3). Compared to other potential fusion partners with collagen-like domain (BTB_c35490, BTB_c38740, Table 3) BTB_c12600 is more similar in its N-terminal collagen-like domain to Pcl7 and contains a peptide that matches perfectly to the consensus motif for incorporation in the surface exosporium. We identified the CTD of BTB_c12600 in *Bt* 407 as the N-terminal 166 amino acids after the last collagen-like repeat unit (Table 4). We replaced this CTD in the chromosome of *Bt* 407 with the 167 amino acid-long CTD of Pcl7 using two consecutive single cross-overs. The construct was verified by sequencing. The resulting recombinant Pcl7-*Bt* strain showed normal growth and sporulation. We induced spore formation and purified spores using standard procedures.

**Table 4.**
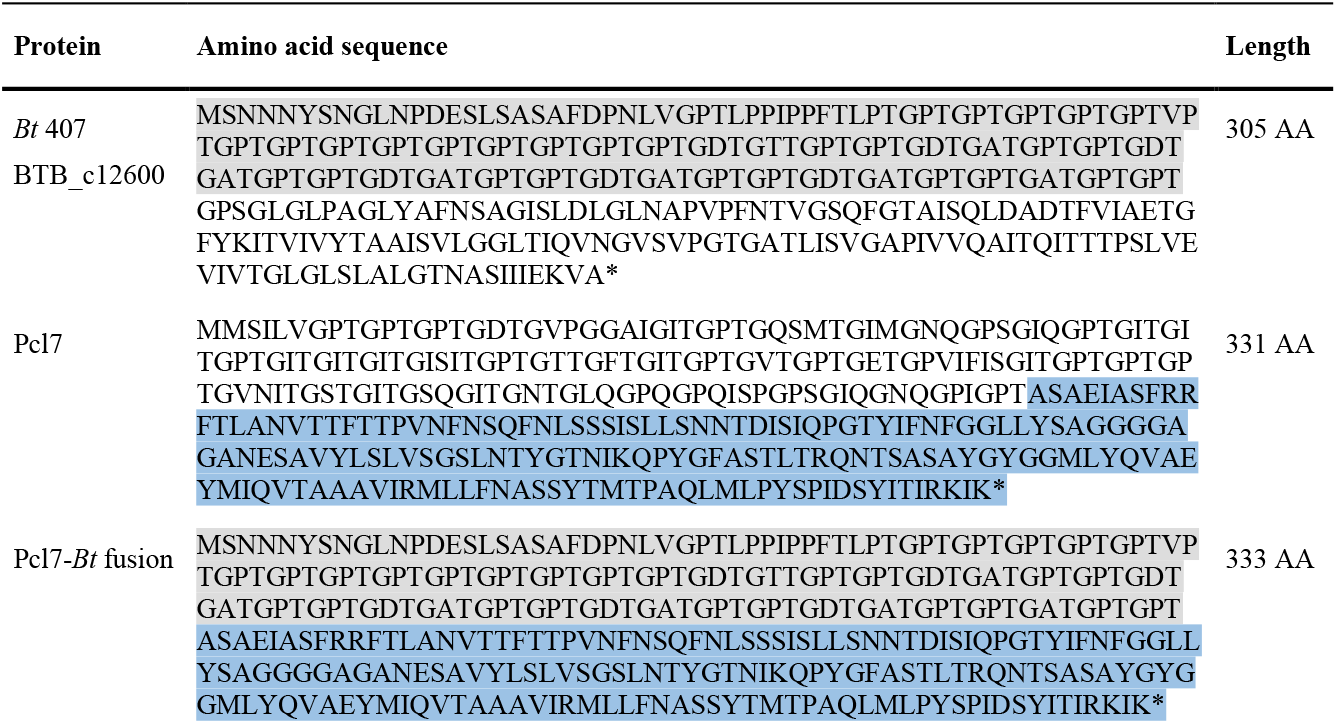
Amino acid sequences used for the Pcl7-*Bt* fusion. Amino acid sequences of the native Bt collagen-like exosporium surface protein BTB_c12600, the Pcl7 protein and the resulting Pcl7-Bt fusion protein. The Bt NTD used in the fusion protein is highlighted in grey, the Pcl7 CTD in light blue. The * represents the stop codon.

To determine adherence of *P. ramosa* and wild type (WT-*Bt*) or *pcl7*-expressing (Pcl7-*Bt*) *B. thuringiensis* spores to *D. magna*, we covalently labelled the bacteria with fluorescein and tracked them in infection assays with different *D. magna* genotypes using fluorescence microscopy (13). C1 *P. ramosa* spores attached to the oesophagus of susceptible *D. magna* (Fig. 2E; 100 % attachment, Fig. 3) but not to resistant hosts (Fig. 3) as previously observed. Vegetative WT-*Bt* and Pcl7-*Bt* cells showed no attachment to the *D. magna* oesophagus but can be observed in the filter setae of both susceptible and resistant *Daphnia* (Fig. 2C), possibly being trapped in the mucus that lines the filter feeding apparatus. WT-*Bt* spores showed attachment neither to the susceptible nor resistant host (0% attachment, Fig. 3), while Pcl7-*Bt* spores attached to the oesophagus of susceptible *D. magna* (Fig 2G). Pcl7-*Bt* attached at low frequency and density to resistant *D. magna* (Fig. 2F; 15 % attachment, Fig. 3), possibly suggesting incorrect glycosylation of Pcl7 (19) in the heterologous *B. thuringiensis* system. Attachment of Pcl7-*Bt* spores to other tissues or known sites of *P. ramosa* attachment such as the hindgut or the external post-abdomen (59) was not observed. Thus, a single domain of collagen-like protein from *P. ramosa* was sufficient to mediate *Pasteuria*-like adhesion properties for recombinant *B. thuringiensis* spores.

**Figure 2.**
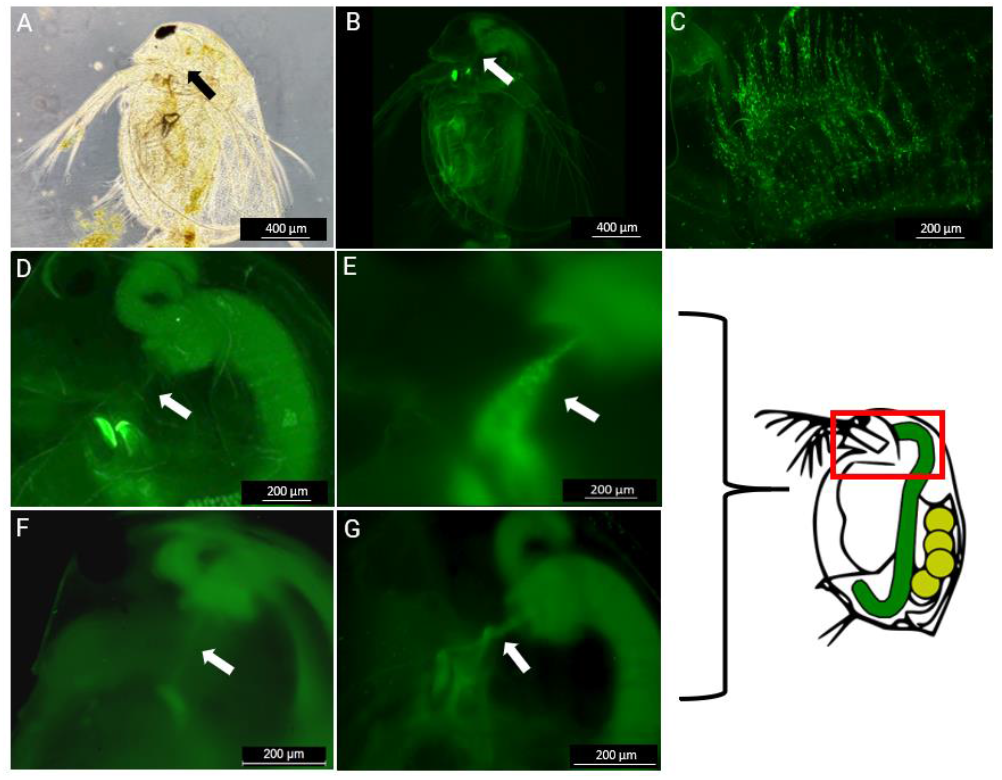
Attachment phenotypes using labelled spores. Microscopy images of the various attachment phenotypes using labelled spores. The red box in the schematic to the right indicates the approximate area (head region) of images D-G. **(A)** Bright field image of a *D. magna* (genotype (=clone) HU-HO-2) showing the entire animal with an arrow indicating the site of the oesophagus. **(B)** Overview of a *D. magna* (genotype HU-HO-2) using a GFP Filter cube showing the entire animal with an arrow indicating the location of the oesophagus. Note the weak autofluorescence of the *D. magna* tissue. **(C)** Labelled *B. thuringiensis* vegetative cells accumulated in the filter setae of a *D. magna* (genotype HU-HO-2). **(D)** Closeup of the *D. magna* foregut with arrow indicating the position of the oesophagus, situated perpendicular to the arrow. The two bright objects left of the arrow are the autofluorescent mandibles. **(E)** Upper body of a susceptible *D. magna* (genotype HU-HO-2) with labelled C1 *P. ramosa* spores attached to the oesophagus (arrow). **(F)** Upper body of a resistant *D. magna* (genotype FI-Xinb3) where labelled Pcl7-*Bt* spores do not aggregate in the oesophagus (arrow). The faint visible light fluorescent band is attributed to autofluorescence. The beginning of the mid gut, visible in the upper right corner, shows fluorescence because labelled spores have been ingested by the *Daphnia*. **(G)** Upper body of a susceptible *D. magna* (genotype HU-HO-2) with labelled Pcl7-*Bt* spores attached to the oesophagus (arrow). The midgut with its appendix (=caecum) is visible on the right.

**Figure 3.**
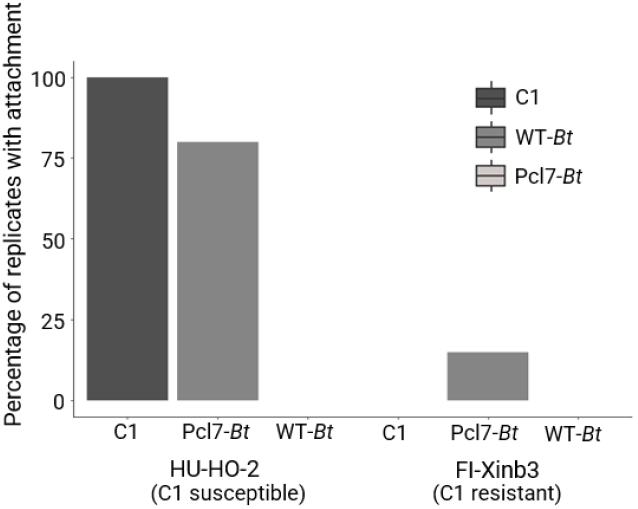
Attachment assay. *D. magna* that are both susceptible (genotype HU-HO-2) and resistant (genotype FI-Xinb3) to C1 *P. ramosa* were exposed to different bacteria: around 5000 labelled C1 *P. ramosa* spores (“C1”), ∼50’000 labelled *B. thuringiensis* WT-*Bt* spores (“WT-*Bt*”), or ∼50’000 labelled Pcl7-*Bt* spores (“Pcl7-*Bt*”). Each isolate was tested on 20 host individuals for each resistotype (N=20, 120 individuals in total).

To determine if Pcl7 presented on *B. thuringiensis* spores can block *P. ramosa* adhesion to *D. magna*, we infected various *D. magna* genotypes with 50 spores of two different *P. ramosa* clones after incubation with a 400-fold excess of the much smaller WT-*Bt* or Pcl7-*Bt* spores. After 5 min exposure, we counted the number of *P. ramosa* spores attaching to the *D. magna* oesophagus. WT-*Bt* had no impact on subsequent *P. ramosa* adhesion for all tested *D. magna* and *P. ramosa* genotypes (Fig. 4). However, Pcl7-*Bt* diminished adhesion of C1 *P. ramosa* spores in four different susceptible *D. magna* genotypes, indicating that Pcl7 was sufficient to block adhesion sites in the host for *P. ramosa*.

**Figure 4.**
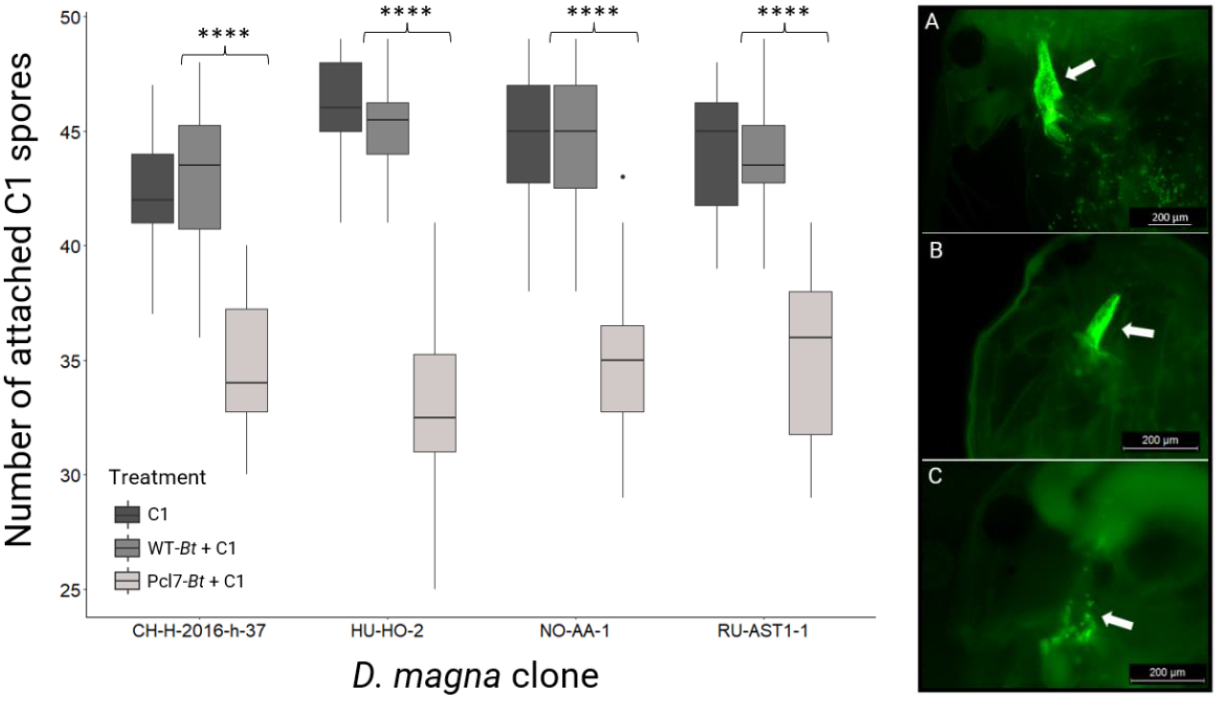
Competitive attachment assay for C1 *P. ramosa*. “C1”– Individuals were exposed to 50 labelled C1 *P. ramosa* spores (picture **A**); “WT-*Bt* + C1” –Individuals were exposed to ∼20’000 *B. thuringiensis* WT-*Bt* spores prior to the addition of 50 C1 *P. ramosa* spores (**B**); “Pcl7-*Bt* + C1” – Individuals were exposed to ∼20’000 Pcl7-*Bt* spores prior to the addition of 50 C1 *P. ramosa* spores (**C**). Each treatment was performed using 20 individual *D. magna*, and the assay was repeated for four different host genotypes (N = 20, 60 individuals per genotype (=clone), 240 individuals in total). A t-test was performed to compare the WT-*Bt* + C1 against Pcl7-*Bt* + C1 means. **** means the t-test is significant with a p-value < 10^−8^. The boxplots display the 25th percentile, median and 75th percentile, while the whiskers display the minimum (Q1-1.5*IQR) and maximum (Q3+1.5*IQR). Outliers are shown as dots. Statistical analysis was done using R (V. 4.1.0, with the R-base package and R packages “ggubr” and “ggplot2”).

We also tested Pcl7-*Bt* against C19 *P. ramosa* that show a different host genotype dependence than C1 *P. ramosa* (the source of Pcl7 in Pcl7-*Bt*). Pcl7-*Bt* had no impact on C19 adhesion to *D. magna* DE-G1-106 (susceptible to *P. ramosa* C19; resistant to C1), but diminished C19 adhesion to *D. magna* HU-HO-2 (susceptible to *P. ramosa* C19 and susceptible to C1) (Fig. 5). Thus, Pcl7-*Bt* could prevent *P. ramosa* adhesion specifically in *D. magna* resistotypes with a Pcl7-receptor (enabling infection by C1), but not in *D. magna* resistotypes without such a receptor. As C19 uses a different receptor than C1, the blocking in HU-HO-2 was probably mediated by steric hindrance between adhering Pcl7-*Bt* and *P. ramosa*.

**Figure 5.**
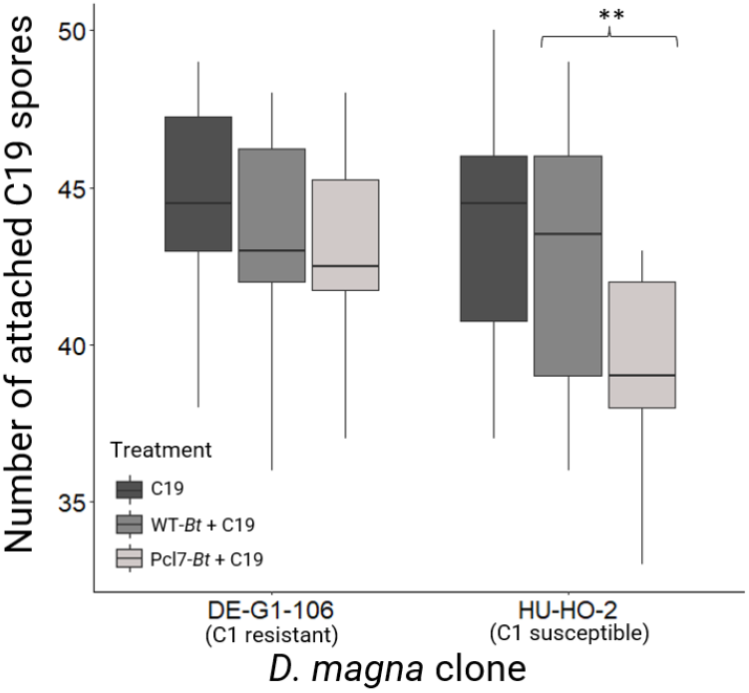
Competitive attachment assay for C19 *P. ramosa*. “C19” – Individuals were exposed to 50 labelled C19 *P. ramosa* spores; “WT-*Bt* + C19” – Individuals were exposed to ∼20’000 WT-*Bt* spores prior to the addition of 50 C19 *P. ramosa* spores; “Pcl7-*Bt* + C19” – Individuals were exposed to ∼20’000 Pcl7-*Bt* spores prior to the addition of 50 C19 *P. ramosa* spores. Each treatment was performed using 20 individual *D. magna*, and the assay was repeated for two different genotypes (N = 20, 60 individuals per genotype, 120 individuals in total). ** *t*-test, p < 0.01.

## Discussion

The mechanism underlying specific coevolution by negative frequency-dependent selection is believed to be driven by the interaction between host and parasite genes. Identifying these genes marks a major step towards understanding this mechanism. In the *Daphnia -Pasteuria* system coevolution is well characterized on a phenotypic level, but the underlying genes are still largely unknown. Here we tested and confirmed the hypothesis that a collagen-like protein (CLP) is crucial for the attachment of the *Pasteuria* parasite to the cuticle of its *Daphnia* host. Polymorphism in the attachment phenotype had previously been shown to be the most important step in the coevolution of the two antagonists (13, 18), and a specific CLP (Pcl7) of *P. ramosa* was linked to the attachment polymorphism (19). Here we used *B. thuringiensis* as a surrogate to express a functional fusion protein harbouring the globular domain of Pcl7. By replacing a single C-terminal sequence in *B. thuringiensis* with the one from *Pasteuria* Pcl7, we were able to create *B. thuringiensis* spores capable of attaching *in vivo* to the host’s oesophagus wall. This result is in line with previous studies that have sought to understand CLP function in bacterial infections using deletion mutants for CLPs (60) or purifying recombinant CLPs (35). Our Pcl7-*Bt* spores not only attached well to four susceptible host genotypes (Figs. 3, 4), but they also attached only sparsely to two resistant host genotypes (Figs. 3, 5), revealing a very similar specificity to that of *P. ramosa* (Fig. 6).

**Figure 6.**
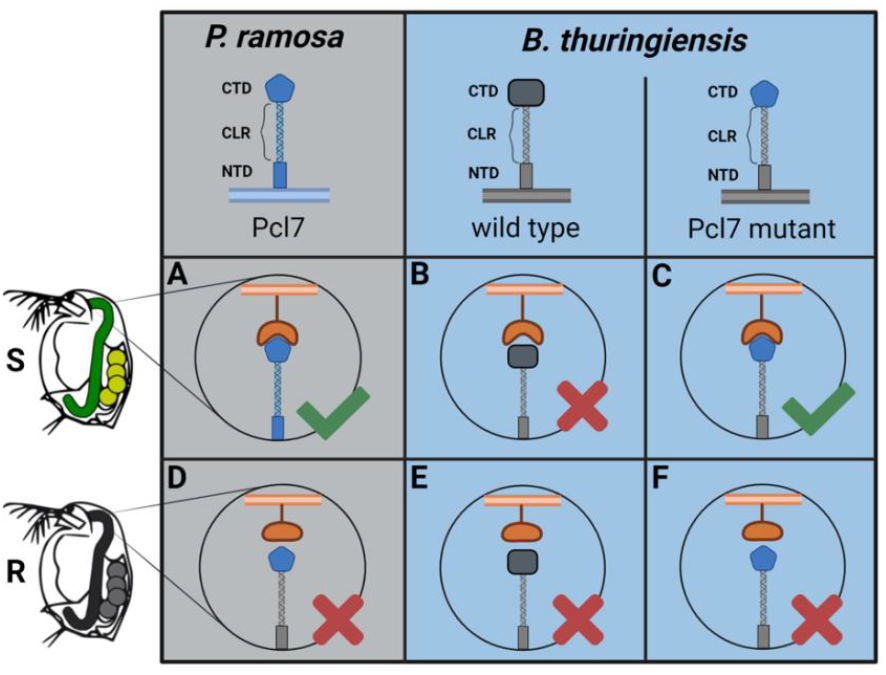
Proposed mechanism for Pcl7-mediated attachment. **(A)** C1 *P. ramosa* ectopically expresses Pcl7 and attaches to the oesophagus of susceptible (S) *D. magna* by binding to host cell surface moieties. **(B)** WT-*Bt* spores do not attach to the oesophagus of susceptible *D. magna* due to the lack of a compatible moiety on the host cell surface. **(C)** Pcl7-*Bt* spores, harbouring the Pcl7 fusion protein attach to the oesophagus of susceptible *D. magna* by binding to the same cell surface moieties as the original Pcl7. **(D)** C1 *P. ramosa* does not attach to the oesophagus of resistant (R) *D. magna*. **(E)** WT-*Bt* spores and **(F)** Pcl7-*Bt* spores do not attach to the oesophagus of resistant *D. magna*. Figure created with Biorender.com

Proteins associated with the spore coat in *B. thuringiensis*, including the BclA homologs that were altered in this study, are predominantly synthesized during sporulation (43). We anticipated that the Pcl7 fusion protein would be present upon completed spore formation. Consistent with this, only spores but not vegetative Pcl7-*Bt* cells adhered to the host’s oesophagus. The specificity of the *B. thuringiensis* spores to a single host tissue thus indicates the functional ectopical expression of the Pcl7 fusion protein, generating the same attachment phenotype as C1 *P. ramosa* spores. This attachment specificity was only seen in hosts susceptible to C1 *P. ramosa*, while in two resistant host genotypes, attachment was either weak or entirely absent. Thus, by replacing part of a single protein, we were able to transform *B. thuringiensis* from non-attaching to attaching, recreating the first step of the infection process. To test if Pcl7-*Bt* spores adhere to the same molecular structure in the oesophagus wall as C1 *P. ramosa* spores we conducted competitive attachment assays. We found that the moieties or potential receptors on the *D. magna* oesophagus cuticle surface are blocked by the Pcl7-*Bt* spores, supporting our hypothesis that the Pcl7-*Bt* interferes with the attachment sites of C1 *P. ramosa*.

The specificity of the Pcl7-*Bt*’s attachment was less pronounced than that of the original *P. ramosa* pathogen, with the difference in attachment between a resistant and a susceptible host genotype being 0 % and 100 % for C1 *P. ramosa*, while it was 15 % and 80 % for the Pcl7-*Bt* spores (Fig. 3). Furthermore, the competitive attachment assays showed that C1 spores were not totally blocked from attachment. Some *Pasteuria* spores were still able to attach after the host had been treated with the Pcl7-*Bt* spores. Finally, to see a visible picture of attachment, we needed much higher spore concentrations of the Pcl7-*Bt*s than the bigger *P. ramosa* pathogen. However, the weak signals observed are certainly caused in part by the rapid germination of *B. thuringiensis* spores once exposed to the *Daphnia*, as germinating spores do not express BclA (43). Furthermore, because *P. ramosa* spores are large (about 5.5 μm diameter) and thus much more visible compared to the *B. thuringiensis* spores (about 1.6 × 0.8 μm) (61), more spores are required to see attachment with the fluorescent microscope.

Other factors, not considered here, may contribute to spore attachment. One such factor may be glycosylation. BclA, and CLPs in general, are known to be highly glycosylated (62, 63). The sequence polymorphism of Pcl7 results in two predicted N-linked glycosylation sites, one of which correlates with the infection phenotype of *Pasteuria* (19) and thus may partly explain the attachment polymorphism. The Pcl7 fusion protein expressed in *B. thuringiensis* may contain different glycans than Pcl7 expressed in *P. ramosa* or might not be glycosylated at all. This might result in altered adhesion properties. Other fusion proteins containing other sequence polymorphisms or glycosylation sites might be used in the future to work out the details of these differences. An analysis of the specific carbohydrate moieties of the Pcl7 glycoprotein could also be done to support this hypothesis (64). However, our data show that such a potential mis-glycosylation had an only a minor impact on adhesion and competition.

Our study verified that the globular part of the Pcl7 protein contributes decisively to the parasite’s ability to attach to the host cuticle, although the molecular mechanisms for the attachment polymorphism are still unclear, as is the composition of the host cuticle surface in the oesophagus. Previous research has shown that CLPs can bind to a variety of molecules and cell surface components such as integrins (28), glycoproteins (34), polysaccharides (65), lipoproteins (35) and mammalian collagen (66). Future studies could focus on identifying the *D. magna* receptor for Pcl7.

## Acknowledgements

We thank J. Hottinger, U. Stiefel, M. Krebs, B. Claudi and members of the Bumann group for help in the laboratory. We thank members of the Ebert group for feedback on the study and the manuscript. We thank S. Zweizig for language editing. We thank Prof. Dr. Ole Andreas Økstad from the Department of Pharmacy of the University of Oslo, Norway for providing the strain and two plasmids used for markerless gene replacement in this study. This work was supported by the Swiss National Science Foundation (SNSF) (grant numbers 310030B_166677, 310030_188887 to D.E.).

## Author contributions

All authors conceived the study. B.H. conducted the laboratory work and analyzed the results. All authors discussed the results. B.H. wrote the manuscript, which was read, edited and approved by all authors.

## Competing interests

All authors declare no competing interests.

